# Endogenous GFP tagging in the diatom *Thalassiosira pseudonana*

**DOI:** 10.1101/2022.09.30.510313

**Authors:** Onyou Nam, Irina Grouneva, Luke C. M. Mackinder

## Abstract

The regulated abundance and spatial distribution of proteins determines cellular structure and function. The discovery of green fluorescent protein (GFP) and fusing it to a target protein to determine subcellular localization revolutionized cell biology. Most localization studies involve introducing additional copies of a target gene genetically fused to GFP and under the control of a constitutive promoter, resulting in the expression of the GFP-fusion protein at non-native levels. Here we have developed a single vector CRISPR/Cas9 guided GFP knock-in strategy in the diatom *Thalassiosira pseudonana*. This enables precise and scarless knock-in of GFP at the endogenous genomic location to create GFP fusion proteins under their native *cis* and *trans* regulatory elements with knock-in efficiencies of over 50%. We show that a previously uncharacterized bestrophin-like protein localizes to the CO_2_-fixing pyrenoid and demonstrate that by measuring GFP fluorescence we can track relative protein abundance in response to environmental change. To enable endogenous tagging, we developed a Golden Gate Molecular Cloning system for the rapid assembly of episomes for transformation into *Thalassiosira pseudonana* via bacterial conjugation. In addition, this versatile toolbox enables CRISPR/Cas9 gene editing, provides a broad range of validated fluorophores and enables future large-scale functional studies in diatoms.

**Significance statement:** Fluorescent protein (FP) tagging is a widely utilized technique for understanding the spatial distribution of proteins. However, introducing extra gene copies under constitutive promoters that randomly integrate into the genome can result in non-biologically relevant expression levels, unwanted genomic mutations and localization artefacts. To overcome this, we developed a novel single vector system capable of CRISPR/Cas9-guided endogenous GFP tagging in a globally important model diatom. This allows scarless GFP knock-in at precise genomic locations resulting in GFP fusions regulated by native promoters/terminators, which facilitates accurate localization and determination of relative protein abundance. Moreover, the developed modular cloning framework is user-friendly and opens the door for high throughput large-scale studies, including FP tagging, knock-out, and knock-in.

## Introduction

Diatoms are a diverse unicellular phytoplankton group that evolved as a consequence of a secondary endosymbiosis event of a red alga (1). They are key marine primary producers responsible for approximately 20% of global net primary production (2) and play central roles in the global carbon and silicate cycles (3–5). In a drive to understand the molecular processes that underpin the ecological success of diatoms, recent years have seen the development of a broad range of genetic tools for two model species, *Phaeodactylum tricornutum* and *Thalassiosira pseudonana*. These include a range of transformation systems such as biolistics, electroporation and bacterial conjugation (6, 7) as well as an array of expression vectors for CRISPR/Cas9 gene editing, heterologous gene expression and fluorescent protein tagging (7–10).

Central to understanding gene function and cellular processes is to know the spatial organization of proteins in the cell. Localizing proteins by tagging the corresponding gene of interest with a fluorescent protein (FP) followed by microscopy to determine the subcellular localization is routinely completed in diatoms (11, 12). However, localization studies often rely on introducing additional copies of the gene under a strong constitutive promoter with these being randomly integrated into the genome. This potentially leads to overexpression of the target gene that might result in incorrect folding and targeting and nonspecific protein-protein interactions (13). In addition, random integration into the genome can result in unpredictable background genotypes and phenotypes (7). An alternative, yet typically more challenging, strategy is to insert a FP seamlessly into the genome in frame with the gene of interest. Known as “endogenous tagging” this approach maintains native promoter and terminator sequences typically resulting in expression levels comparable to native levels (reviewed (13)). In model organisms where homology-directed recombination (HDR) is dominant (e.g. yeast) endogenous tagging has enabled large-scale proteome tagging studies (14). However, in many eukaryotes a double stranded break is required for efficient HDR. This has been shown to be the case for diatoms where targeted genomic integration via HDR only occurs if (i) a double-stranded break has been introduced by a targeted endonuclease (15, 16) or (ii) components of the non-homologous end joining pathway have been knocked down (17). Previous reports have indicated a higher occurrence of HDR in *T. pseudonana* compared to *P. tricornutum* (15–18). The now routine application of CRISPR/Cas9 gene editing in diatoms (7, 19) opens up the potential for endogenous tagging. CRISPR/Cas engineering was first demonstrated in 2016 (8, 20) and can be achieved via stable integration into the genome of Cas9 and sgRNA (20), editing using RNP complexes (21) or expression of Cas9 and sgRNA(s) from an episome (9).

In this study we develop a seamless CRISPR/Cas9 guided knock-in strategy that enables endogenous GFP-tagging of proteins expressed from their native genomic locus and without the co-integration of an antibiotic marker. To achieve this, we first built a Golden Gate based modular cloning (MoClo) framework for episomal delivery via conjugation in *T. pseudonana*. This framework enables rapid and scalable cloning for a broad range of applications and requires no specialist equipment for transformation. We demonstrate versatility by validating a range of promoter-terminator pairs, testing a broad spectrum of fluorophores, and performing CRISPR/Cas9 gene editing. We then expand the methodology to enable precise genomic knock-in of GFP for endogenous protein tagging using a single vector system.

## Results

### Development of a Modular Cloning framework in *T. pseudonana*

To enable the versatile application of fluorescent protein tagging and CRISPR/Cas9 gene editing combined with the ease of bacterial conjugation we developed an episomal based Golden Gate MoClo framework (Fig. 1A; see SI appendix Table S1 for vectors). Golden Gate cloning uses Type IIS restriction enzymes that cut outside of their recognition sites allowing user defined overhangs (syntax). Assemblies are hierarchical, typically individual level 0 parts (i.e. promoter, coding sequence, FP, terminator) are assembled into level 1 transcriptional units, that are further assembled into a level 2 transformation vector. To be compatible with previously developed MoClo parts (8) and maintain some compatibility with the universal Loop (uLoop) assembly framework (10) we used the Plant/*Chlamydomonas* MoClo syntax (22, 23). Our standard level 2 episome used for diatom transformations contains: an episome maintenance part, *CEN6-ARSH4-HIS3*, that prevents random integration into the genome and enables it to replicate and be maintained as an episome inside *T. pseudonana* (6); the origin of Transfer (*oriT*) required for vector transfer during conjugation (6) and; the nourseothricin N-acetyl transferase (NAT) gene conferring resistance to the antibiotic nourseothricin. As the *CEN6-ARSH4-HIS3* and *oriT* are always required in combination these were combined to form a single level 1 part and occupy position 1 in our L2 assemblies with the *NAT* marker typically position 2. Depending on application, positions 3-5 of our L2 assemblies can be occupied with different L1 plasmids assembled from L0 parts or custom made with the correct flanking syntax e.g. sgRNAs (Fig. 1A).

**Fig. 1.**
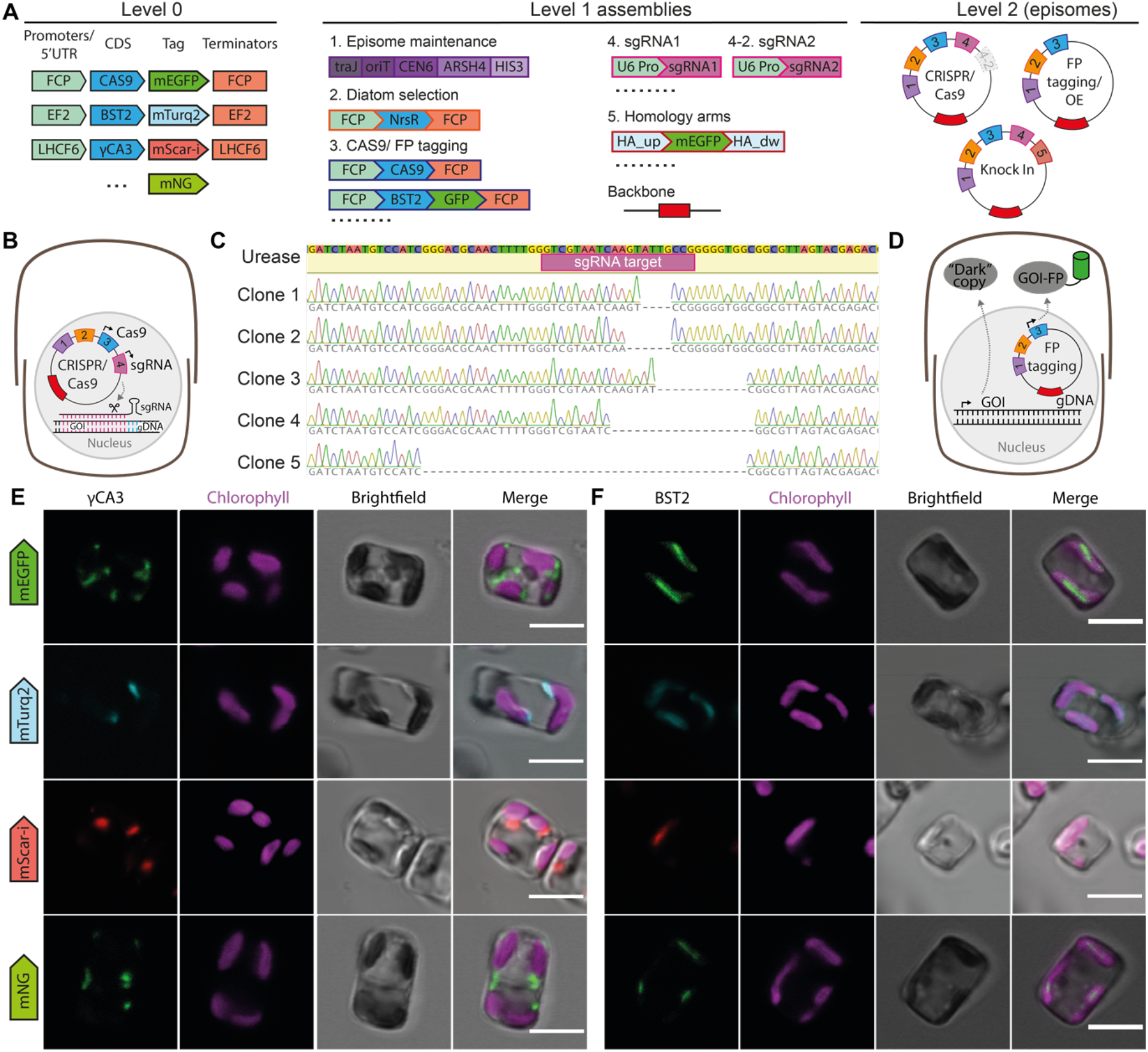
Establishment of a Golden Gate based cloning framework for episome delivery of constructs in *T. pseudonana*. *(A)* Overview of Golden Gate assembly parts developed. These parts can be combined to assembly episomes for CRISPR/Cas9 gene editing, fluorescent protein (FP) tagging and FP knock-in. *(B)* Cas9 and sgRNA targeting the Urease gene were expressed from an assembled episome. *(C)* After CRISPR/Cas9 gene editing the Urease locus was PCR amplified, cloned and sequenced from episome containing colonies. *(D)* Episomes expressing a gene of interest (GOI) C-terminally fused to a fluorescent protein were assembled and transformed. This results in a tagged version and the native untagged (“dark”) version of the protein. *(E)* and *(F)* Confocal imaging of developed fluorophores validated on *γ*CA3 and BST2. Scale bars: 5 μm.

We initially tested our assembly framework by assembling an episome for CRISPR/Cas9 editing of the urease gene using previously validated sgRNAs (8) (Fig. 1B). After transformation and selection of *T. pseudonana* colonies the urease locus was PCR amplified and cloned. Sequencing indicated that we could efficiently edit the urease gene using our episome assembly and delivery framework (Fig. 1C). We next tested our Golden Gate episome system for fluorescent protein tagging by expressing an additional copy of our gene of interest (GOI) C-terminally fused to monomeric enhanced GFP (mEGFP; GFP from here on; Fig. 1D). We first validated this on the previously localized (24) mitochondrial gamma carbonic anhydrase 3 (*γ*CA3) using the fucoxanthin chlorophyll a/c-binding protein (FCP) promoter and FCP terminator (Fig. 1E). We next applied it to a previously unlocalized target gene, bestrophin-like protein (THAPSDRAFT_4820), which we termed BST2. BST2 has previously been shown to be transcriptionally and translationally induced at low CO_2_ levels (25, 26). Bestrophin-like proteins have been shown to play an important role in the CO_2_ concentrating mechanism of *Chlamydomonas reinhardtii*. They are proposed to channel HCO_3_^−^ into the thylakoids that traverse the Rubisco containing pyrenoid, enabling its sequential dehydration to CO_2_ and fixation by Rubisco (27). We showed that BST2-GFP localized to a defined elongated spot at the center of the chloroplast, and we tentatively assign this to the membranes that traverse the pyrenoid (Fig. 1F). To expand the available fluorophores and regulatory regions of our system we tested and validated the fluorophores mNeonGreen, mScarlet-i and mTurquoise2 (Fig. 1E and F) and the previously developed but untested promoter/terminator pairs of elongation factor 2 (EF2) and light harvesting complex f 6 (LHCF6) from the uLoop system (10) (SI appendix Fig. S1).

### Development of a scarless endogenous GFP tagging strategy

The “gold standard” of fluorescent protein tagging is to scarlessly insert a fluorescent protein into the genome resulting in the target protein being labelled whilst still under the control of its native *cis* and *trans* regulatory elements (13). To achieve this, we set out to establish a CRISPR/Cas9 guided HDR GFP knock-in approach that could be delivered on a single episome (Fig. 2A). To test our approach, we attempted to endogenously GFP tag BST2, which we previously localized via expression as a fusion protein directly from episomes (Fig. 1F).

**Fig. 2.**
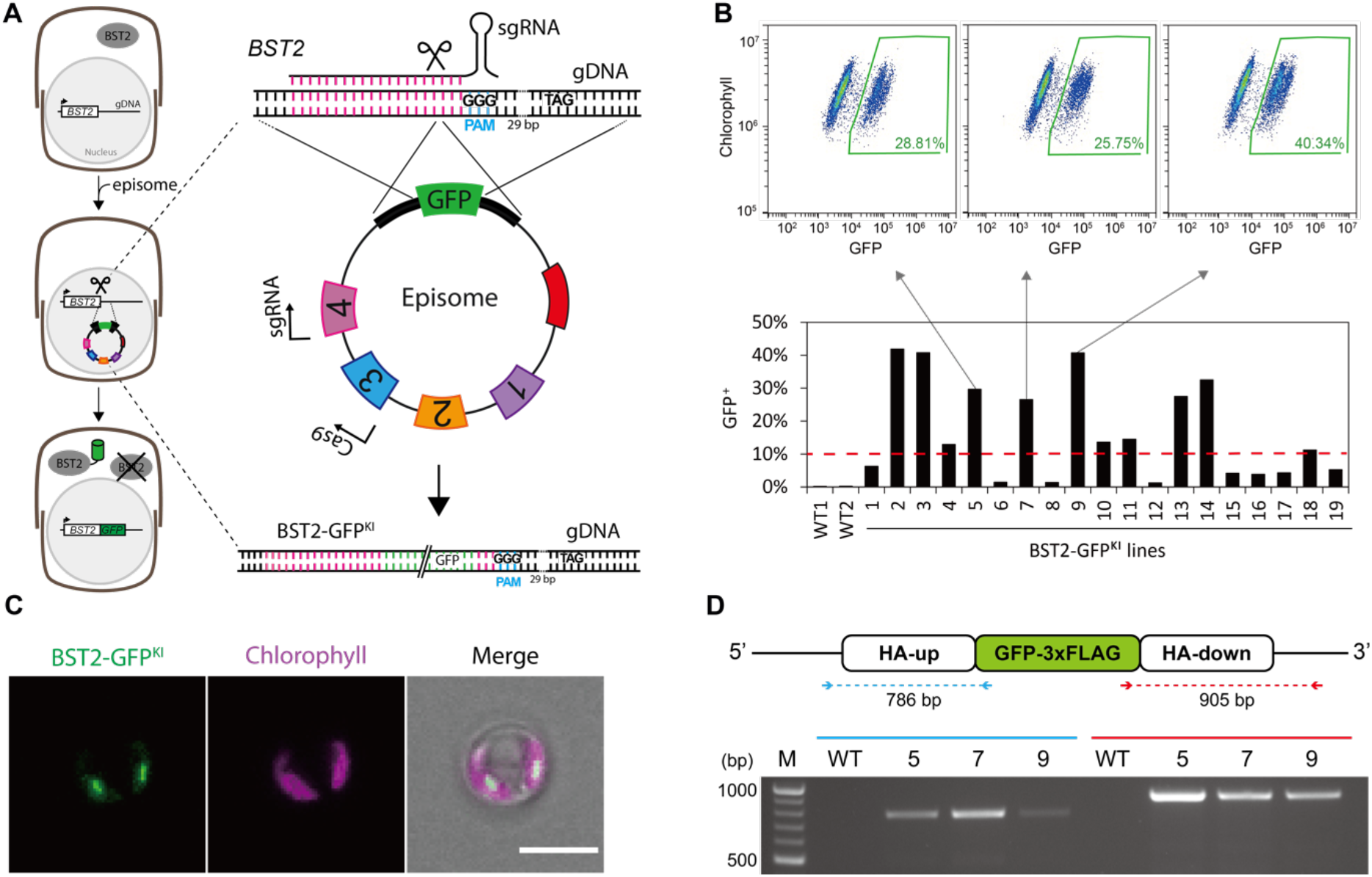
Endogenous GFP tagging strategy overview and transformant screening. *(A)* Endogenous GFP tagging strategy. *T. pseudonana* is transformed with an episome containing (1) *CEN6-ARSH4-HIS3* and *oriT*, (2) *NAT* selectable marker, (3) Cas9, (4) sgRNA and the repair template containing GFP. Cas9 cutting enables homology-directed repair by the repair template generating a GFP gene fusion. To prevent Cas9 re-cutting after homology directed repair the repair template disrupts the sgRNA recognition region. The layout of the schematics and indicated base pairs represent what was used for the BST2-GFP^KI^. *(B)* Screening colonies for fluorescence: Nourseothricin resistant colonies were picked, grown in liquid, and then screened for fluorescence using a CytoFLEX flow cytometer. The green box represents the percentage of GFP positive cells. Lines with >10% (dashed red line) of GFP positive cells were classified as successful BST2-GFP^KI^ lines. *(C)* Confocal images for a BST2-GFP^KI^ line identified from flow cytometry. Scale bar: 5 μm.(D) Genotyping of BST2-GFP^KI^ lines showing the presence of GFP at the targeted genomic locus of BST2.

Using CRISPick (28) (see methods) we selected a sgRNA with a predicted on-target score of 0.7 that cut 40 bp upstream of the BST2 stop codon. We designed homology arms that covered both sides of the predicted Cas9 cut site (Fig. 2A). In this case successful BST2-GFP knock-in (BST2-GFP^KI^) would lead to the disruption of the sgRNA target region preventing re-cutting by Cas9. GFP knock-in would result in the GFP being inserted 13 amino acids upstream from the stop codon (Fig. 2A). AlphaFold modeling (29) of BST2 and BST2-GFP^KI^ indicated that this would be in a region of low predicted structure and unlikely to impact BST2 folding (SI appendix Fig. S2). Homology arms were either PCR amplified from genomic DNA or custom synthesized to enable domestication (removal of BsaI and BpiI restriction sites) when required. The domestication of any homology arms was done via synonymous mutations to avoid changes at the amino acid level. Homology arms were assembled into a L1 plasmid separated by the in-frame GFP sequence. The L0 GFP vector used contained no stop codon, with the stop codon coming from the native stop codon within the downstream homology arm. The resulting homology arm GFP flanked L1 construct (GFP repair template) was assembled into an L2 plasmid. This L2 plasmid contained five L1 parts including [1] *CEN6-ARSH4-HIS3* and *oriT*, [2] *NAT* selectable marker, [3] Cas9, [4] sgRNA and [5] GFP repair template (Fig. 2A). Assembled episomes were transformed into *E. coli* EPI300 cells carrying the pTA_Mob mobility plasmid that enables transfer of the episome to the target organism. Episome and pTA_Mob containing *E. coli* were incubated with *T. pseudonana* and selected on nourseothricin containing plates.

### Identifying and genotyping endogenously tagged GFP knock-in lines

Nourseothricin resistant colonies were transferred to 24-well plates for growth in liquid media. Liquid grown transformants were screened via flow cytometry using a CytoFLEX that enables rapid screening in a 96-well plate format. Initial data for our BST2-GFP^KI^ construct indicated that 58% (11/19) of transformants were GFP positive when a stringent 10% cut-off for GFP positive cells was used (Fig. 2B and SI appendix Fig. S3). Lines that showed a high population of GFP cells were selected for microscopy (Fig. 2C) with the localization of BST2-GFP^KI^ showing a linear traversion at the center of the chloroplast most likely within the thylakoid membrane traversing the pyrenoid. This data corresponded to when BST2-GFP was expressed from an episome (Fig. 1F), indicating that in this case neither the native expression levels nor insertion of GFP 13 amino acids upstream of the C-terminal affected localization. We confirmed scarless insertion of *GFP* via amplification and sequencing of the insertion flanking regions using oligos located outside the homology arms to exclude the possibility of amplicons originating from the episome (Fig. 2D). To ensure that Cas9 mediated DNA cutting was required for GFP insertion, and to check that GFP could not be expressed from the episome prior to HDR, we assembled an episome that lacked the sgRNA targeting *BST2*. Screening of 15 colonies via flow cytometry showed no GFP positive colonies (SI appendix Fig. S4A). The presence of the episome was confirmed by PCR against the *NAT* gene (SI appendix Fig. S4B). This indicates that Cas9 DNA cleavage is required for efficient GFP knock-in and GFP is not expressed without successful integration into the genome.

### Determining knock-in efficiency and isolating lines with close to 100% GFP positive cells

From our flow cytometry data (Fig. 2B) and GFP imaging we typically saw a mixed population of GFP positive and GFP negative cells suggesting the presence of mosaic colonies, where different genotypes are present within the same colony (30). To isolate pure clonal lines, we developed a flow cytometry fluorescence cell counting pipeline and optimized this for the GFP knock-in of *γ*CA3 (Fig. 3A and B). Our pipeline typically involves three rounds of flow cytometry. Transformants are picked and grown in the presence of antibiotics in 24-well plates with weekly subculturing. After 2-3 weeks (~6-9 generations) these are then screened via flow cytometry. Lines with the highest percentage of GFP positive cells are further propagated for ~3-4 generations and re-screened under antibiotic pressure. Once lines are close to 100% GFP positive they are transferred to antibiotic free media to drive episome loss. First round flow cytometry data gave us the percentage transformants with positive GFP signal. From this data we conclude that knock-in efficiency of at least one allele being successfully tagged in *T. pseudonana* for BST2 is 58% (Fig. 2B) and for *γ*CA3 is 81% (Fig. 3B). Further rounds of flow cytometry screening indicated that multiple lines with close to 100% GFP positive cells could be isolated using this approach (Fig. 3C). Confocal microscopy of these lines showed a typical mitochondrial localization pattern (Fig. 3D).

**Fig. 3.**
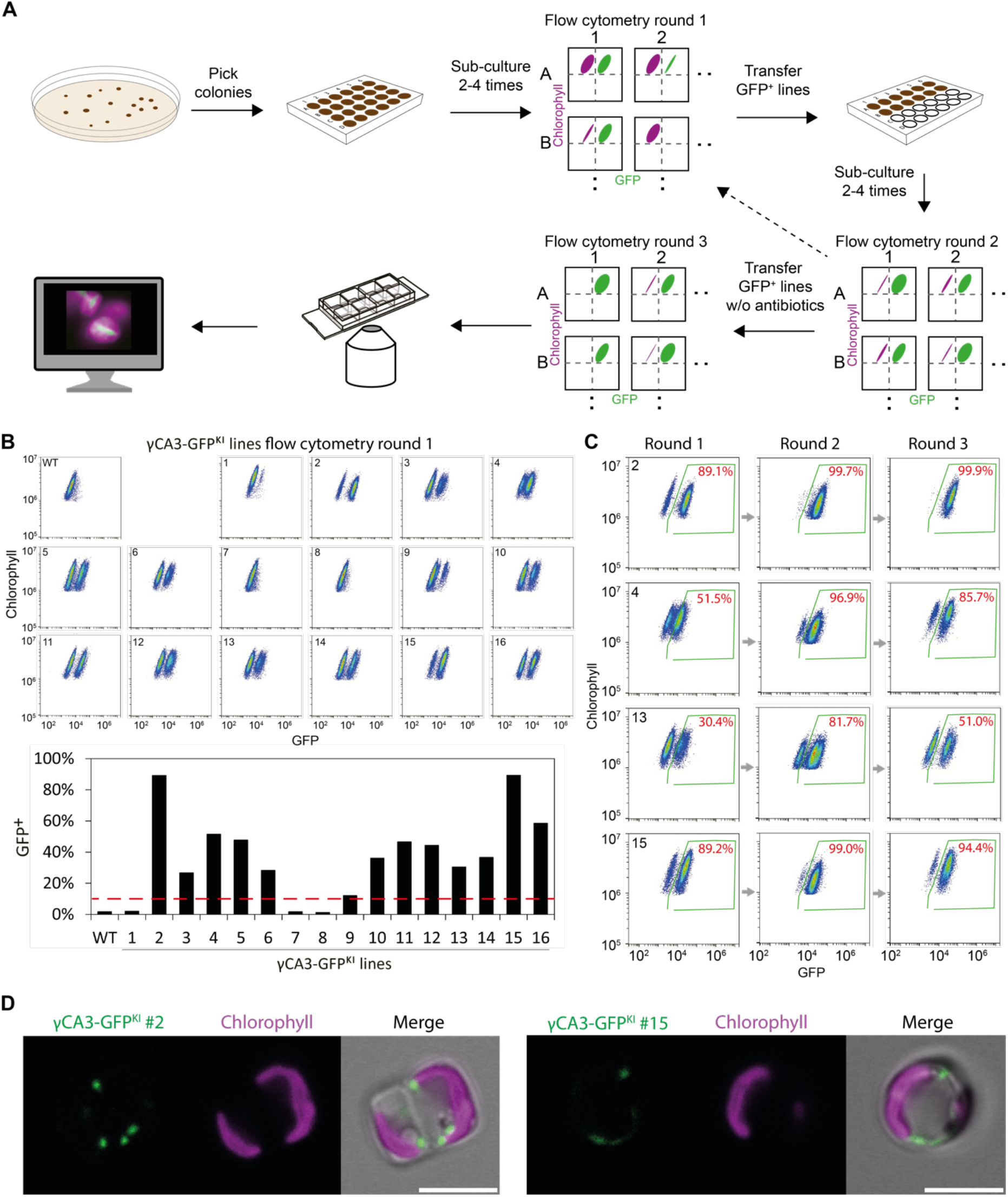
Development of a flow cytometry screening pipeline. *(A)* Developed pipeline to identify and select colonies with high GFP knock-in fluorescence. *(B)* Round 1 flow cytometry counting of *γ*CA3 transformants. Bar chart corresponds to the above plots. Lines with >10% (dashed red line) of GFP positive cells were classified as successful *γ*CA3-GFP^KI^ lines. *(C)* Four lines from round 1 flow cytometry counting were subcultured on antibiotics for ~3-4 generations and the percentage of GFP positive cells counted (Round 2). Lines were transferred to antibiotic free media for episome curing, subcultured for ~3-4 generations and then counted (Round 3). *(D)* Confocal microscopy of two *γ*CA3-GFP^KI^ lines. Scale bar 5 μm.

### Screening enables biallelic editing

Vegetative *T. pseudonana* is diploid. GFP knock-in at only a single allele potentially is not stable due to chromosomal cross-over (31) that could result in GFP removal and revertant back to a wild-type genotype. In addition, in a hetero-allelic knock-in GFP fluorescence would not represent native protein abundance due to presence of an untagged copy. To examine if GFP has been knocked into both alleles in our clonal lines we PCR amplified the knock-in genomic locus with primers that flank the homology arms. In a biallelic knock-in we would expect a single band corresponding to the locus containing GFP, whereas in a non-clonal or mono-allelic line we would expect an additional lower molecular weight wild-type band. In both BST2-GFP^KI^ and *γ*CA3-GFP^KI^ lines with close to 100% GFP positive cells we could detect only a single band indicating successful GFP integration into both alleles (Fig. 4A and B). Sequencing across the homology arm and genomic DNA junctions and across the GFP homology arm junctions indicates scarless GFP insertion (Fig. 4C and D). Immunoblotting against the GFP tag of the confirmed biallelic lines indicated that both BST2-GFP^KI^ and *γ*CA3-GFP^KI^ were at the expected size (Fig. 4E and F).

**Fig. 4.**
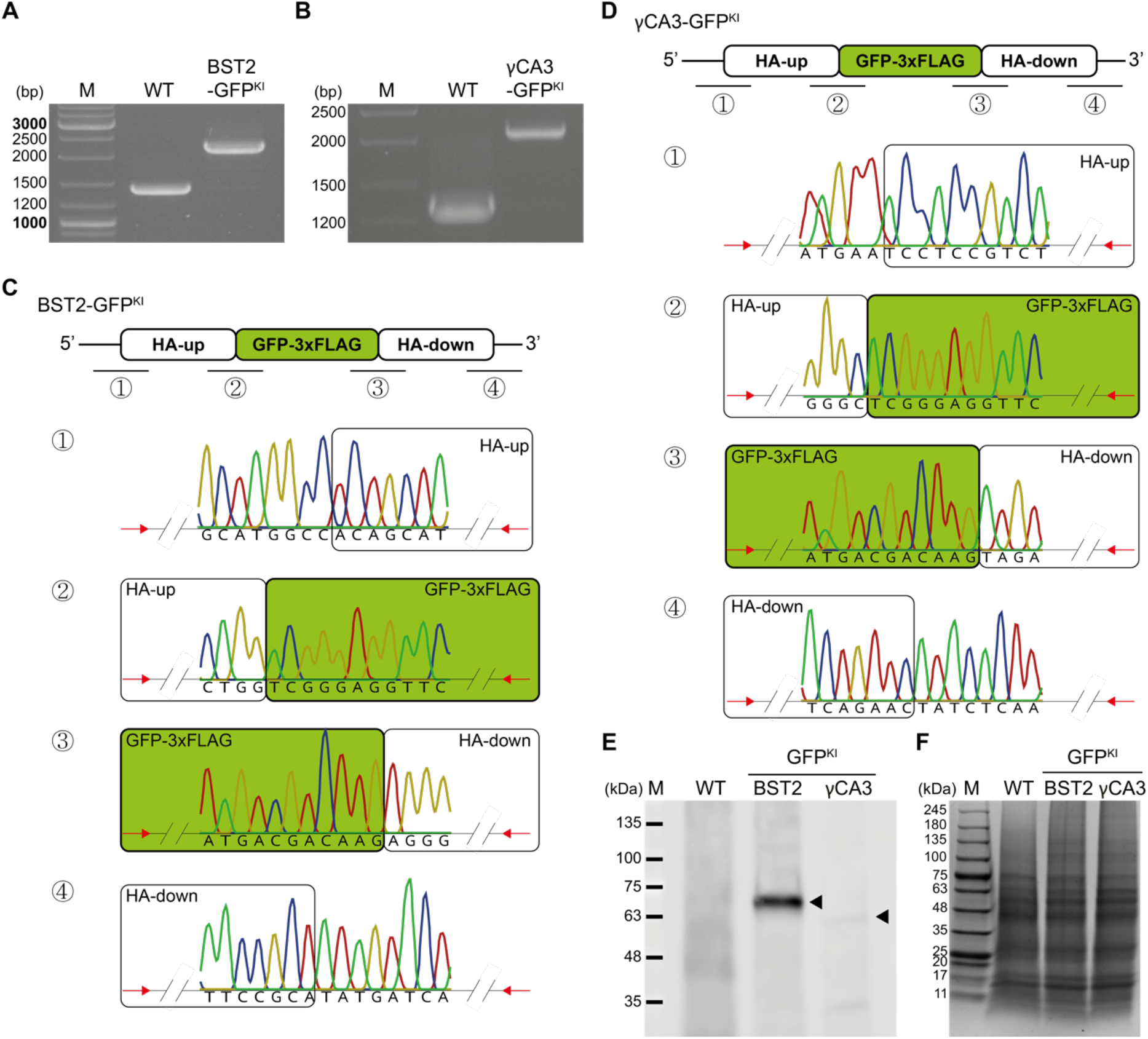
Confirmation of biallelic GFP knock-ins. *(A)* and *(B)* PCR to confirm biallelic editing of clonal lines. *(C)* and *(D)* Sequencing of junctions confirming scarless integration of GFP into the genome at the target location. *(E)* Immunoblot against GFP in BST2-GFP^KI^ and *γ*CA3-GFP^KI^. Arrow heads indicate the expected sizes of BST2-GFP (73 kDa) and *γ*CA3-GFP (57 kDa). *(F)* Coomassie stained SDS-PAGE loading control.

### Relative protein abundance can be tracked by fluorescence intensity

As GFP knock-in lines are under their native regulatory elements we tested if changes in relative protein abundance could be tracked in response to an environmental change. As BST2 was previously shown to increase in abundance ~4-fold at low CO_2_ (26) we grew a BST2-GFP^KI^ line at both high and low CO_2_ and then measured GFP intensity per cell using flow cytometry (Fig. 5A). We saw an ~2-fold increase in GFP fluorescence in low CO_2_ grown cells demonstrating that relative protein abundance can be tracked using our GFP knock-in lines (Fig 5B and C).

**Fig. 5.**
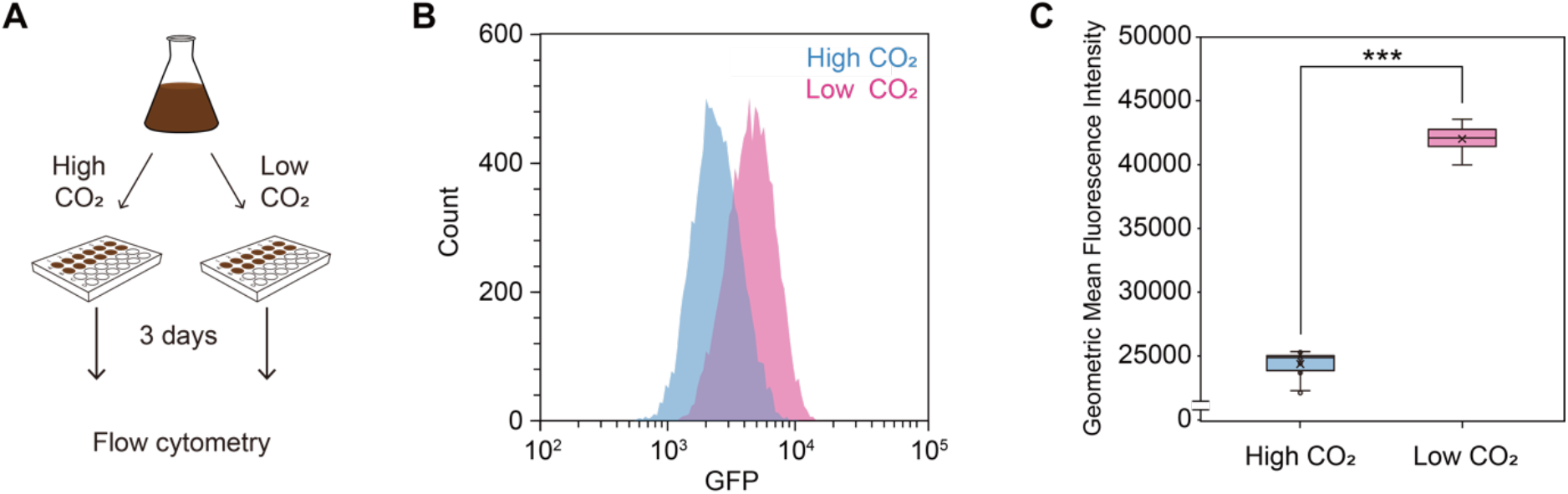
Relative protein abundance can be tracked in GFP knock-in lines. *(A)* A biallelic BST2-GFP^KI^ line was grown in ambient CO_2_, transferred to both high CO_2_ and low CO_2_ for three days then GFP intensity measured by flow cytometry. *(B)* and *(C)* Low CO_2_ grown cells show a clear shift in fluorescence intensity (***p<0.001, Students *t* test).

## Discussion

We developed a single plasmid-based approach for high efficiency endogenous GFP tagging in the diatom *T. pseudonana*. Our method enables the scarless insertion of GFP at a precise genomic locus to generate a fluorescent protein fusion without the co-integration of a selection marker. This ensures that native *cis* and *trans* regulatory regions are maintained enabling relative protein abundance to be tracked in response to perturbations. To demonstrate the power of our approach we applied it to BST2, a bestrophin-like protein. The localization of endogenously GFP tagged BST2 to an elongated central chloroplast region corresponds to the thylakoids traversing the pyrenoid. Tracking BST2-GFP^KI^ relative abundance using flow-cytometry showed an increase at low CO_2_ supporting previous transcriptional (25) and proteomic (26) data. As all of the native protein copies are GFP fusions in our biallelic BST2-GFP^KI^ lines, we envision that single molecule counting approaches can be used in the future to determine absolute protein numbers (32). Collectively, the data supports an analogous role of BST2 to *Chlamydomonas* bestrophin-like proteins implicated in the CO_2_ concentrating mechanism (27). In *Chlamydomonas*, BST1-3 are low CO_2_ upregulated and localized to the pyrenoid periphery with a proposed role to deliver HCO_3_^−^ from the stroma to the thylakoid lumen where it is subsequently dehydrated by a carbonic anhydrase to release CO_2_ within the pyrenoid for fixation by Rubisco (27). Further evidence supporting an analogous mechanism in diatoms is the presence of a θ-carbonic anhydrase in the lumen of pyrenoid-traversing thylakoids in *P. tricornutum* that is required for efficient CO_2_ fixation (33). As algal bestrophin-like proteins are further characterized it will be interesting to see if they play a central role in evolutionary diverse pyrenoid based CO_2_ concentrating mechanisms.

Whilst HDR endogenous FP knock-in approaches have been developed for several model systems, the seamless knock-in of FPs has yet to be achieved for eukaryotic algae. In photosynthetic eukaryotes a GFP knock-in strategy has been established in the model plant *Arabidopsis thaliana* by sequential expression of a sgRNA and GFP donor template in a parental line expressing Cas9 (34). In-line with our single plasmid approach, the authors also tested the presence of all components on a single T-DNA construct, however this failed to result in heritable GFP knock-in (34). As *T. pseudonana* is diploid, an additional challenge associated with endogenous tagging is the knock-in of the GFP at both alleles. To overcome this, we maintained transformants under antibiotic selection pressure and developed a rapid and scalable flow cytometry screening pipeline. This resulted in the generation of homoplasmic knock-in lines.

During the establishment of our GFP knock-in approach we only tested homology arms of ~600 bp and the knock-in of GFP (~800 bp). Further tests to identify the minimum homology arm size requirement could lower homology arm synthesis costs in the future. In addition, exploring the limitations of knock-in fragment size would be useful to determine if the approach could be used for multi-gene/pathway knock-in at neutral genomic sites for biotechnology applications. We also only tested guide block knock-ins where the GFP is inserted into the Cas9 cut-site. It has been demonstrated in other organisms that HDR can be offset from the cut-site which would then enable GFP to be inserted directly prior to the stop codon and increase sgRNA flexibility. However, knock-in efficiency can rapidly drop off as the cut-site to HDR-site distance increases (35).

Our knock-in efficiency of 58-81% appears to be high compared to other systems (34, 36, 37). This reduces the effort required for screening and selection of successful GFP knock-in lines. One potential explanation for this high efficiency is the presence of all the necessary engineering elements on a single episome that can be continually maintained through antibiotic selection. The high efficiency of our system indicates that the same framework could be used for insertions, substitutions and functional knock-outs at specific target sites. However, for biallelic editing additional screening strategies, other than fluorescence-based flow cytometry, would need to be established.

To enable the successful and easy application of our system we first developed and validated a Golden Gate based episome assembly framework for multi-purpose genetic engineering in *T. pseudonana* (Fig. 1). This included the episomal testing of two additional promoter/terminator pairs, the validation of three additional fluorophores (mTurquoise2, mScarlet-i and mNeonGreen) and the establishment of episomal based CRISPR/Cas9 gene editing. The Golden Gate based episomal assembly framework allows large, multi-part constructs to be rapidly and easily assembled. Episomal delivery by bacterial conjugation enables the delivery of large constructs, avoids random integration into the genome and requires no specialized equipment such as a Gene gun. This engineering framework now opens the door for large-scale fluorescent protein tagging and mutant library generation in diatoms. Similar studies in other systems have proven to be extremely powerful for understanding gene function and biological processes (14, 38–41).

## Methods

### Strains and growth conditions

*Thalassiosira pseudonana* CCAP1085/12 (equivalent strain CCMP1335) was obtained from the Scottish Culture Collection of Algae and Protozoa (CCAP) and grown axenically at 20°C with a continuous illumination of ~50 μmol photons m^−2^ s^−1^. in F/2 medium (42, 43) prepared with artificial sea water (33 g L^−1^, Instant Ocean SS15-10).

### Episome assemblies using Golden Gate cloning

Level 0 (L0), Level 1 (L1) and Level 2 (L2) plasmids were assembled by Golden Gate (GG) cloning (44) using a range of custom parts, uLoop (10) parts and Golden Gate MoClo Plant Kit parts (45) (SI appendix Table S1). The assembly framework and syntax was according to Patron et al. 2015 (22). Available *T. pseudonana* expression cassettes for Cas9, the *NAT* selection marker, *U6* promoter and sgRNA backbone were used to construct L1 vectors (8). With both Cas9 and the *NAT* selection marker under the expression of the *FCP* promoter and terminator. The yeast element *CEN6-ARSH4-HIS3* (6) was domesticated for GG cloning by removing BsaI and BpiI restriction sites and cloning it into a L1 plasmid (pICH47732) together with oriT and traJ. The resulting L1 part permits low-copy episomal replication in diatoms without random integration into the genome (6). Depending on application (i.e. gene knock-out, FP-tagging or GFP knock-in) 3-5 L1 parts can be assembled into an L2 plasmid (termed episome) for transformation into *T. pseudonana*.

Each one-tube, one-step, restriction-ligation GG assembly contained 40 fmols of each component. Final reaction volumes of 20 μl were incubated in 10x ligase buffer (Promega) with 10 units restriction enzyme (BsaI-HFv2, NEB or BpiI, Thermo Fisher Scientific) and 10 units T4 DNA ligase (HC, 20 units μl^−1^, Promega). In a thermocycler, the reaction was switched between 16°C and 37°C at 5 min intervals for 30 cycles, followed by 5 min at 37°C, then increased to 65°C for 20 min. 2.5 μl were transformed into 50 μl DH5α *E.coli* cells and selected on antibiotic containing plates.

### *Transformation of* T. pseudonana

Golden Gate assembled episomes were transformed into electrocompetent *E.coli* (TransforMax EPI300) already carrying the mobility plasmid pTA_Mob (46) (gift from R. Lale). *E.coli* colonies carrying both mobility and cargo plasmid were used for subsequent conjugation. Bacterial conjugation was performed according to (6) with the minor adjustment that post conjugation *T. pseudonana* cells were left to recover under standard growth conditions (20°C, 50 μmole photons m^−2^ s^−1^) for 24 hours instead of four hours before spreading onto selective media containing 100 μg ml^−1^ Nourseothricin.

### Gene targets and homology arm design

The following genes were targeted for GFP knock-in: BST2 (THAPSDRAFT_4820) and γCA3 (THAPSDRAFT_265081).

A template for HDR was integrated into an episome as a further L1 component. It consisted of i) a domesticated 600 bp sequence upstream of the stop codon of the targeted gene (HA_up), ii) in-frame GFP without a stop codon and iii) a domesticated 600 bp sequence downstream of but including the stop codon (HA_down). sgRNAs used in this experiment were selected so that the cut site they induced would be as close to the stop codon as possible. For both genes a guide blocking approach was used whereas after insertion of the GFP via HDR the sgRNA recognition site would be split preventing further cutting. Domesticated 600 bp homology arms (upstream and downstream of the selected Cas9 cut site) containing restriction sites and junctions for GG cloning were either gene synthesized (Twist Bioscience) or amplified from genomic DNA. GFP was fitted with a short flexible linker and a 3xFLAG tag.

### Genotyping/screening for HDR

Fifty microliters were harvested from each transformant line grown in liquid selection medium. Pellets were resuspended in 20 μl of lysis buffer (20 mM Tris-HCl pH 8.0, 10 mM EDTA, 10% Triton X-100) and kept on ice for 10-20 min, followed by incubating at 95°C for 10 min. Cell debris was removed by brief centrifugation and 1 μl of supernatant used as PCR template. The OneTaq Hot Start Quick-load 2x Master Mix with GC buffer (M0485L, New England BioLabs) was used for PCR. All primers used in this study are listed in SI appendix Table S2.

### Confocal microscopy

Images were taken with a laser scanning microscope (LSM880, Zeiss) using 63x objective with a 1.4 numerical aperture. Excitation lasers and emission filters were as follows: mTurquoise2 excitation 458 nm, emission 463 - 535 nm; GFP and mNeonGreen excitation 488 nm, emission 481 - 541 nm; mScarlet-i excitation 561 nm, emission 561 - 633 nm. For chlorophyll excitation we used the same excitation as the corresponding fluorophore with emission 642 - 712 nm.

### Flow cytometry

GFP expression was analyzed by flow cytometry using the CytoFLEX LX355 (Beckman Coulter) analyzer. Forward scattered (FSC) and side scattered photons by the 488 nm laser were used to determine diatoms from culture debris. FSC-height versus FSC-area signal was used to isolate single events from sample aggregates. Chlorophyll autofluorescence excited by 561 nm laser and emitted photons detected with 675/25 filter was used to ensure all samples were fully intact. GFP fluorescence excited by the 488 nm laser was detected by avalanche photo diode detector with band pass filter 525/40. CytExpert software (Beckman Coulter) was used for analysis.

### SDS-PAGE and immunoblotting

Cells were harvested by centrifugation (3,000 *g*, 10 min). Pellets were resuspended in cell lysis buffer (20 mM Tris-HCl pH 8.0, 50 mM NaCl, 0.1 mM EDTA, 12.5% glycerol, 1x protease inhibitor cocktail [Pierce]) and sonicated for 1 min for cell disruption. The protein extracts were separated by SDS-PAGE using 10% Protean TGX, 50 ul wells gel (BioRad). Subsequently, gels were stained with coomassie blue or transferred to PVDF membrane for immunoblotting. The GFP protein was probed with an anti-GFP antibody at 1:1,000 dilution (G10362, Thermofisher) and visualized using a goat anti-rabbit IgG (H+L) cross-adsorbed secondary antibody, Alexa Flour 488 (Invitrogen). The fluorescence was imaged using Typhoon-5 (Amersham).

### Vector and strain distribution

Upon publication all developed vectors will be made freely available via Addgene, and all developed strains available from the authors.

## Acknowledgements

The funding was supported through a United Kingdom Research and Innovation Future Leaders Fellowship to L.C.M.M (MR/T020679/1), Biotechnology and Biological Sciences Research Council Grants (BB/T017589/1, BB/S015337/1, BB/R001014/1) and Engineering and Physical Research Council Grant (EP/W024063/1). The authors would like to thank the University of York Biosciences Technology Facility for confocal microscopy and flow cytometry equipment access and support.

## Author Contributions

L.C.M.M. and I.G. conceived the knock-in strategy. I.G., O.N. and L.C.M.M. designed the experiments and wrote the manuscript. O.N. and I.G. carried out the experiments.

## Supplementary Figures

**Fig. S1.**
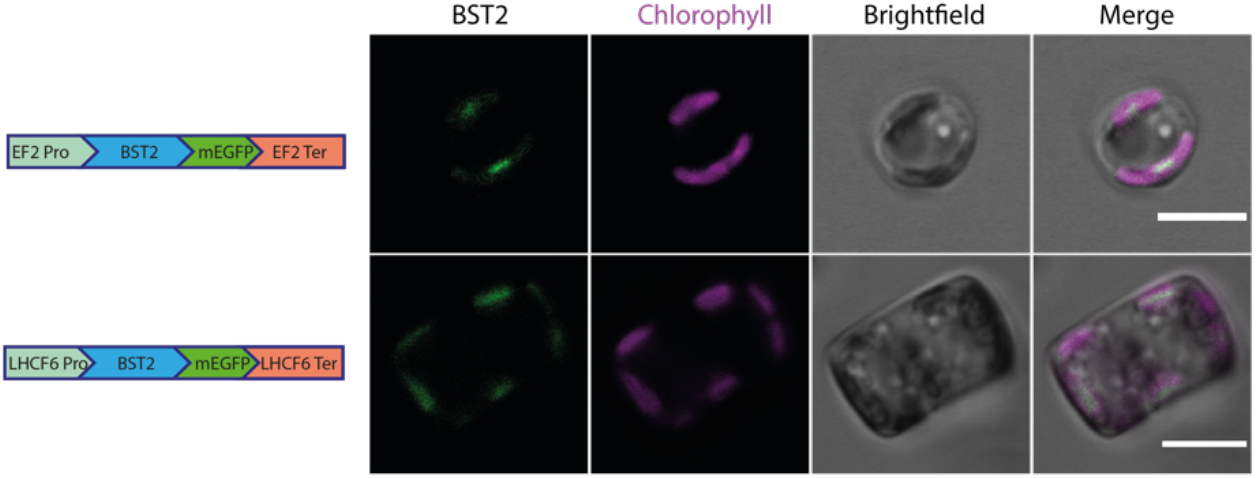
Validation of two additional promoter and terminator pairs. The elongation factor 2 (EF2) promoter and terminator and the light harvesting complex f 6 (LHCF6) promoter and terminator were validated for fusion protein expression in *T. pseudonana*. Scale bar: 5 μm.

**Fig. S2.**
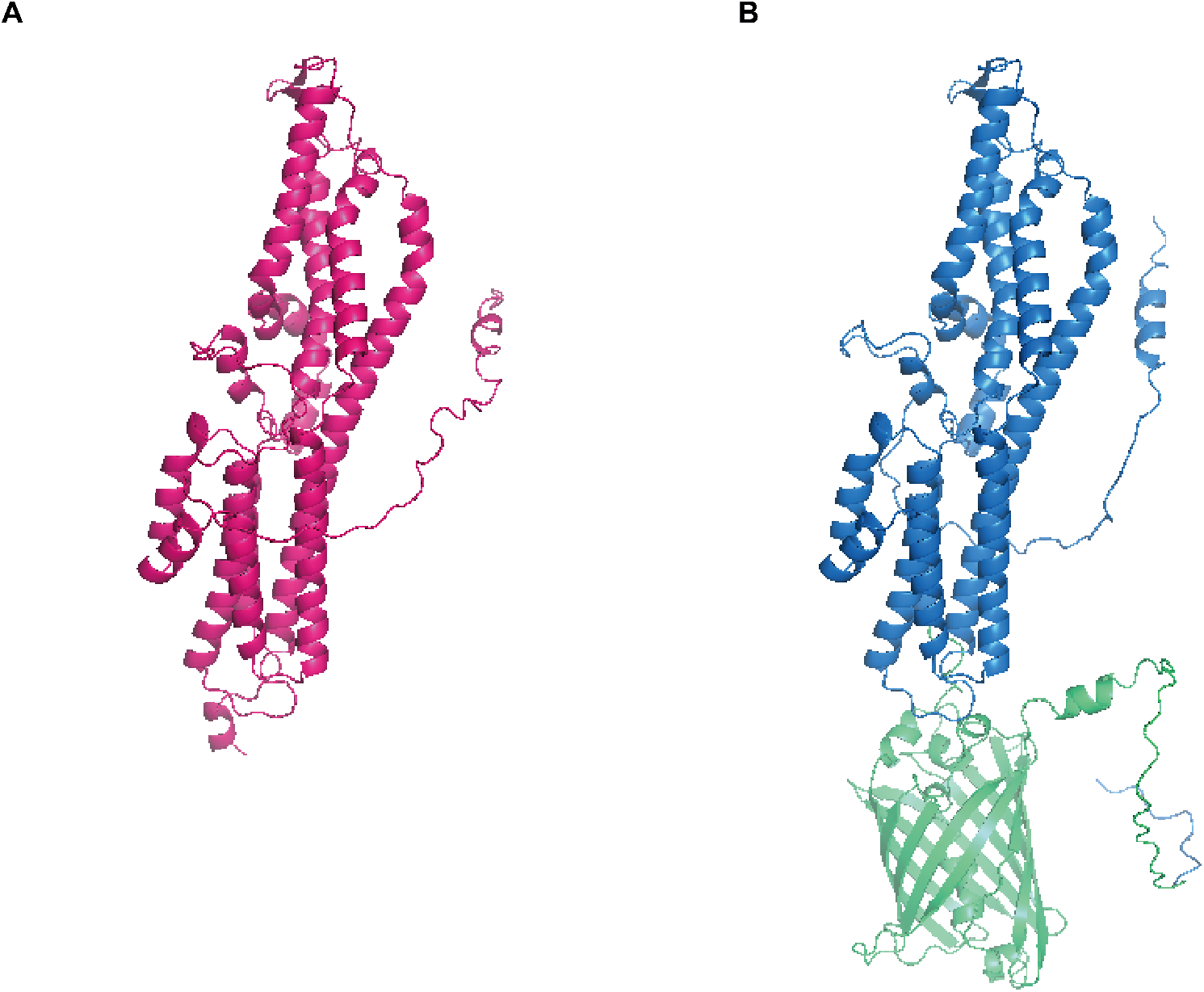
AlphaFold model of BST2 and BST2-GFP^KI^. *(A)* AlphaFold model of BST2. *(B)* AlphaFold model of BST2-GFP^KI^ showing that GFP knock-in has minimal predicted structural changes. Blue: BST2, Green: mEGFP-3xFLAG.

**Fig. S3.**
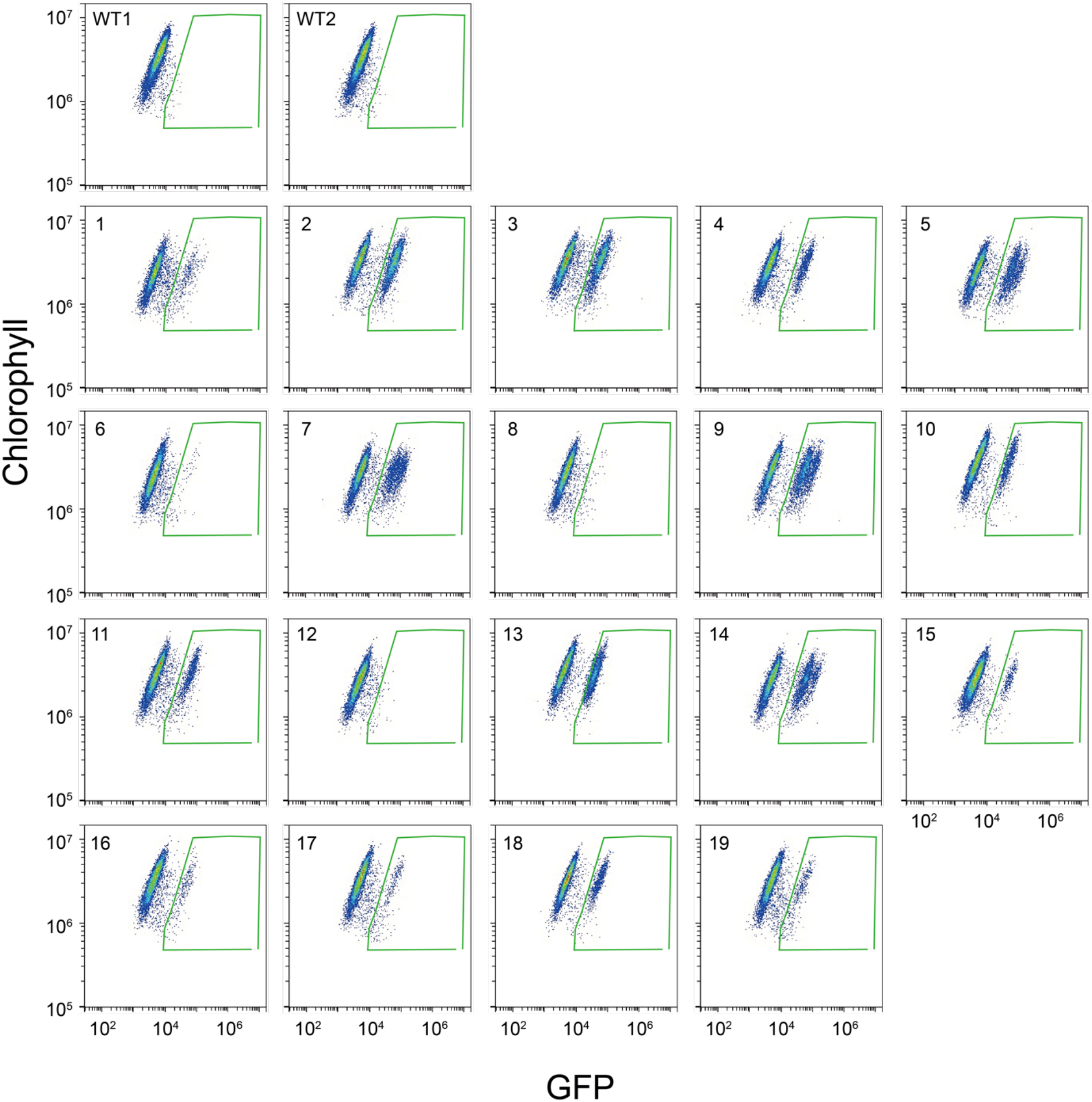
Flow Cytometry screening of BST2-GFP^KI^ lines. Round 1 flow cytometry counting of BST GFP knock-in transformants. Green boxes are gates used for counting GFP positive cells.

**Fig. S4.**
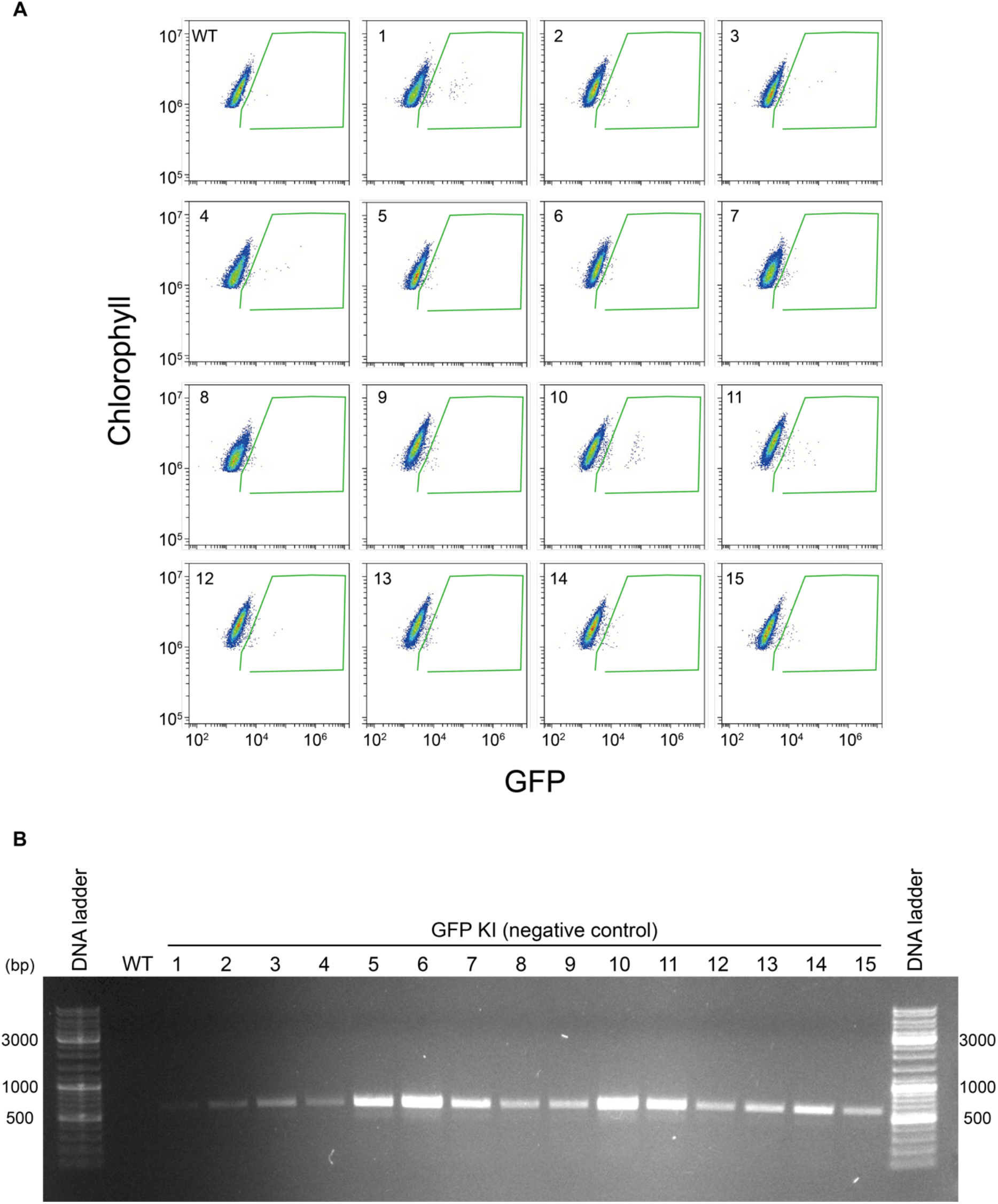
The sgRNA is required for GFP knock-in and fluorescence. *(A)* Flow cytometry screening of 15 colonies picked after transformation with a GFP knock-in episome containing homology arms for BST2 GFP knock-in but that lacked the sgRNA. No GFP positive colonies were identified. *(B)* Colony PCR against the *NAT* gene in the episome backbone to confirm the presence of the episome in the flow cytometry screened colonies.

